# Contribution of cortical layers to human dynamic functional connectivity

**DOI:** 10.1101/2023.04.11.536354

**Authors:** Patricia Pais-Roldán, Seong Dae Yun, Shukti Ramkiran, Tanja Veselinović, Irene Neuner, Markus Zimmermann, Jörg Felder, N. Jon Shah

## Abstract

Transitory states of the brain, sometimes linked to particular mental disorders, can be identified by conducting a dynamic analysis of connectivity in functional magnetic resonance imaging (fMRI) data. Here, we investigated whether the connectivity between cortical *laminae* (in contrast to between cortical *areas*), can vary during the course of a resting-state scan and whether this could impact the identification of brain states in the healthy brain.

We performed a sliding-window connectivity analysis of high-resolution fMRI data and found that the differential participation of the cortical layers in the overall cortical connectivity constitutes a more dynamic feature of the brain compared to region-to-region connectivity, with two main states (superficial vs. deep-laminae connectivity) fluctuating over time. Laminar connectivity restricted to the default mode network appeared relatively stable (superficial connectivity preferred), while the central executive and salience networks showed a fluctuating laminar preference. We anticipate that the dynamic analysis of connectivity focused on the different depths of the cerebral cortex could play an important role in characterizing healthy brain states as well as in identifying novel targets of psychiatric diseases.

## INTRODUCTION

Most neurons fire relatively often in a healthy brain and, as a result, continuous wave-shaped signals are observed in functional recordings of the healthy living brain (Berger, 1929, Buzsaki and Draguhn, 2004). Whole-brain imaging methods, e.g., functional magnetic resonance imaging (fMRI), allow brain function to be assessed at a network level, typically in the form of temporal correlations, which can serve not only to characterize the healthy brain but also to distinguish patients with neuropsychiatric diseases from healthy controls (Menon, 2011, Canario, 2021).

Most functional connectivity studies are based on scan-specific measures (i.e., one connectivity matrix per scan), however, mental states cannot be explained exclusively by single connectivity patterns. Instead, subjects typically transit between two or more mental states, and hence, static-connectivity measures may be an over-simplistic interpretation of reality (Chang and Glover, 2010, Allen et al., 2014, Hutchison et al., 2013). In order to detect fluctuations in brain states, the brain imaging community has reformulated the common connectivity analysis to include finer time evaluations, e.g., using a sliding-window scheme by which functional connections are assessed every few seconds within 1-2 minute blocks, or a variety of other comparable methods (Lurie et al., 2020, Fu et al., 2019, Zhou et al., 2019, Warnick et al., 2018, Shappell et al., 2019, Chakravarty et al., 2019). This allows periods of stronger or weaker connectivity between areas to be identified that would otherwise be missed in a static analysis. The connectivity patterns, their variability, and duration associated with the time-varying states have been the focus of numerous recent publications assessing differences between healthy subjects and diverse groups of patients (Niu et al., 2019, Jiao et al., 2021, Pang et al., 2020, Zhang et al., 2018, Ahmadi et al., 2021, Naro et al., 2018, Cao et al., 2019, Gu et al., 2020, Rabany et al., 2019, Wu et al., 2021, Fallahi et al., 2021, Fu et al., 2020, Zhu et al., 2019, Wei et al., 2020, Kaiser et al., 2016, Wong et al., 2021, Zhang et al., 2021, Demertzi et al., 2019, Ramkiran, 2022), demonstrating the clinical relevance of tracking such connectivity changes in the human brain. Moreover, the reconfiguration of these dynamic connectivity patterns upon stimulation further suggests that functional dynamic connectivity analysis may be of value in novel therapeutic approaches (Grami et al., 2021).

To date, largely owing to a lack of methods to image the whole brain with high spatial resolution and sufficient coverage, the literature has focused on region-of-interest (ROI)-based network analysis (static or dynamic), disregarding the depth at which the connections take place in the cortex. Recently, novel high-resolution fMRI sequences have allowed us not only to detect the connections between remote functionally-different areas but to do so with sub-millimetre resolution, enabling the evaluation of resting-state networks at multiple cortical depths (Huber et al., 2020, Yun et al., 2019, Yun and Shah, 2017, Yun et al., 2020, Yun and al., 2021, Pais-Roldán et al., 2020, Pais-Roldán et al., 2023, Yun et al., 2022). For instance, we have previously shown that, in healthy subjects, areas of the default mode network (DMN) are strongly correlated in the superficial layers, while the central executive control network (CEN) involves connections between deeper cortical territories (Pais-Roldán et al., 2023).

Whether the preferred cortical depth at which brain areas communicate can change during the course of an fMRI scan remains unexplored. In this context, the question arises whether high-resolution fMRI can be used to provide added value to investigate dynamic connectivity states. If so, the newly incorporated dimension (cortical depth) could radically improve our understanding of particular neuropsychological diseases. In this study, we used laminar-fMRI to assess the brain connectivity dynamics of the resting state in healthy subjects. We investigated the effect of using connectivity matrices with different levels of spatial content to determine brain states, characterized the depth-dependent dynamic connectivity states, and evaluated the differences in state identification for three resting-state networks (the triple network, i.e., DMN, CEN and salience network (SN)).

## RESULTS

### Relevance of the cortical depth for tracking brain-state transitions

The k-means algorithm clusters the connectivity matrices from different time points in a way that the within-group variance is minimized. We hypothesized that the use of input matrices based on laminar connectivity instead of ROI connectivity could alter the results of the clustering process. The relevance of the different levels of connectivity across the cortical depth in brain state determination (see **Fig. 1**) was demonstrated with a higher number of state switches observed when applying k-means to the laminar connectivity matrices compared to the typical ROI-based analysis (conditions 25 and 26, see **Fig. 2a** –state switches of all conditions and subjects-, **Fig. 2b** –mean number of state switches per subject- or **Fig. 2c** –mean number of state switches per condition-). We provide a complete description of all conditions tested and their individual outcomes in **Supplementary Text**. Compared to any other scenario, the analyses of connectivity purely based on layer information (conditions 25 and 26 in Fig. 2, i.e., where the inputs were either the total level of connectivity at each one of six cortical depths or a layer-to-layer connectivity matrix) resulted in the highest level of dynamicity (highest unrest index). It is important to note that subject motion did not correlate with the number of state switches of any condition. Our results suggest that the dynamics occurring along the cortical depth, independently of ROI localization, could play an important role in the transition between brain states.

**Figure 1.**
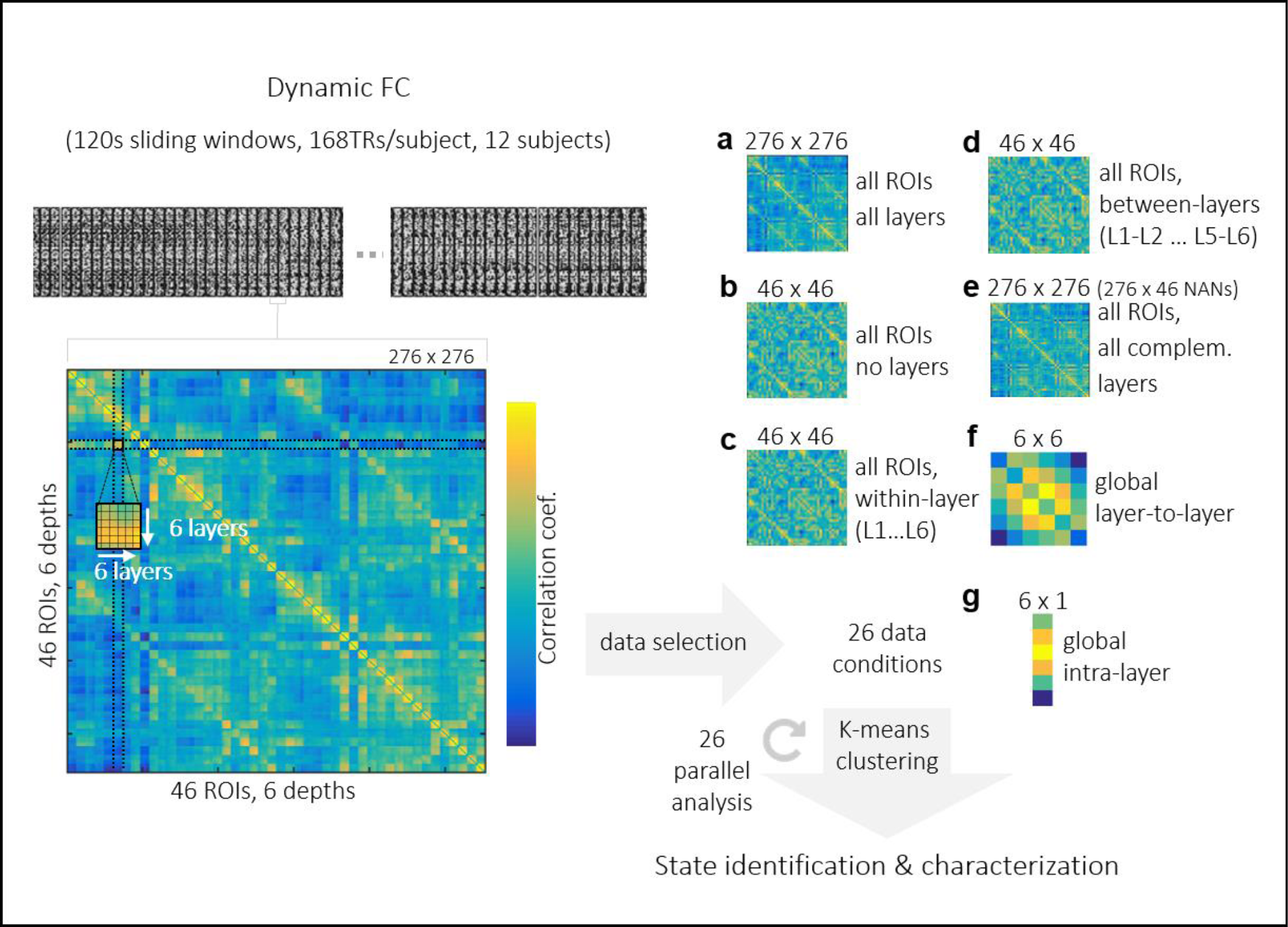
Overview of the main analysis steps to assess the dependence of brain state fluctuation on input data. The original resolution of a functional connectivity matrix was 276, or six cortical depths of 23 ROIs in two cortical hemispheres. Functional connectivity was calculated in sliding windows over concatenated rs-fMRI scans from twelve volunteers (time dimension was 168 TRs × 12 subjects). To test the influence of the information contained in the available data (“conditions”, e.g. a number of ROIs and/or layers) on brain state identification, the original connectivity matrix was fed to the dynamic connectivity analysis either (**a**) in its entirely or (**b**) modified by averaging across layers (leaving only ROI information), (**c**) by discarding all data except for those relevant to one particular cortical depth (this matrix was generated for each of the six depths), (**d**) by considering only the connection between specific pairs of layers (15 possible combinations), (**e**) by excluding within-layer connections (i.e., leaving only information in complementary layers), (**f**) by averaging across ROIs to focus on the layer-to-layer global connectivity (disregarding the spatial coordinates on the brain surface) or (**g**) by further averaging across layers to obtain a proxy of degree centrality for each of 6 cortical depths. The number pairs “m x n” over each matrix indicate the size of the input matrix. In each of the 26 conditions, a k-means algorithm was used to split the corresponding time-varying matrices into coherent groups, presumably representing different brain states.

**Figure 2.**
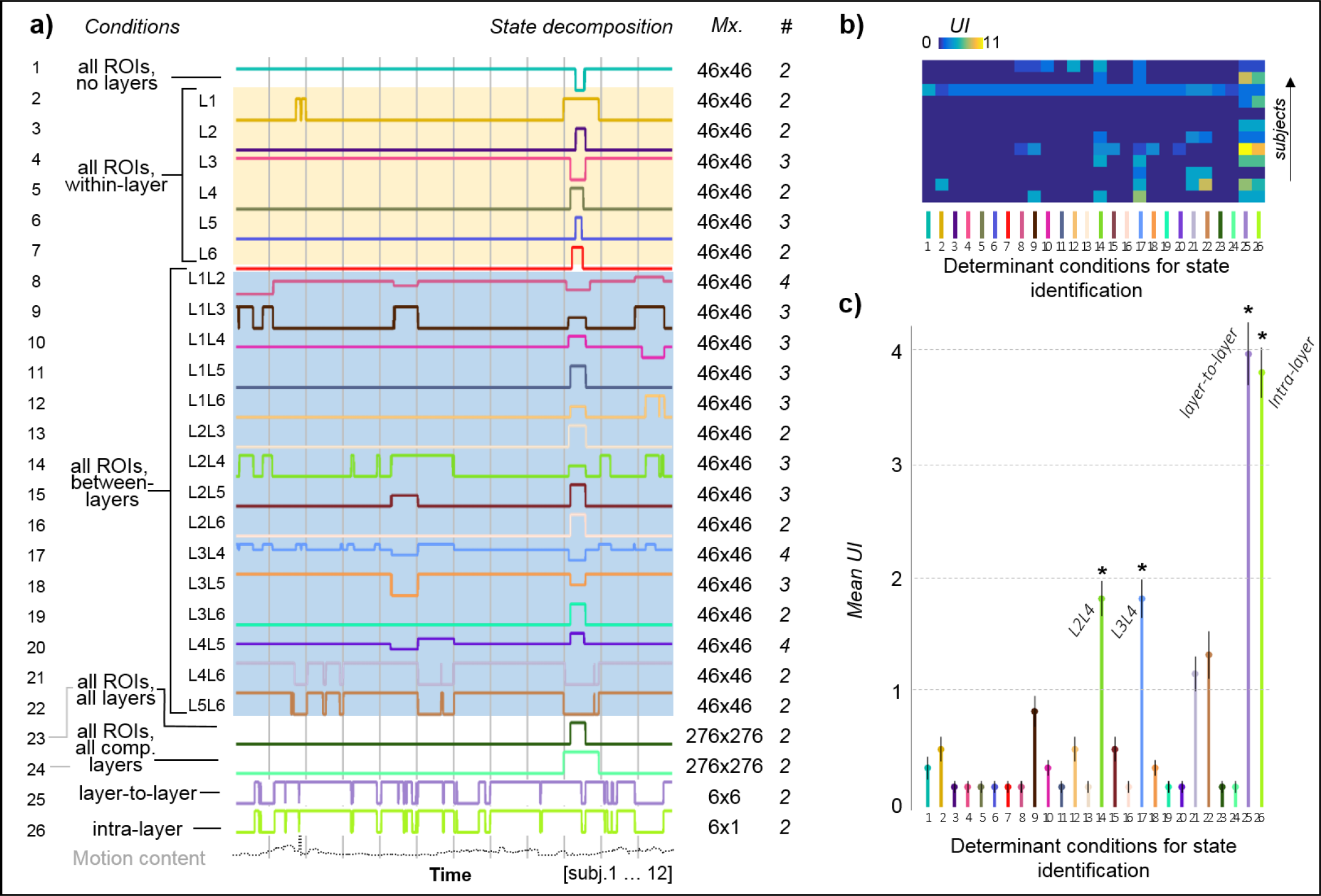
Dynamicity of the brain state associated with input data. (**a**) Brain states were identified based on the dynamic connectivity patterns of 26 sets of data (“conditions”, as presented in Fig. 1 –mind the different order, the “all ROIs and layers” case (“a” in Fig. 1) is condition 23 here-) and the equivalent vector of the motion content. Vertical grey lines represent subject separation. The numbers on the right indicate the size of the matrix (ignoring time dimension) that was subjected to dynamic correlation analysis (“Mx”) and the number of different states detected per condition (“#”). (**b**) Unrest index (UI) computed as the number of brain state switches (colour-coded) for each subject (y-axis), for each of the different analyses (x-axis), following the same order from left to right as in “a” from top to bottom). (**c**) Mean unrest index (UI) ± SE, calculated per condition (n=12 subjects). An asterisk (*) highlights conditions that showed a UI significantly different (p<0.05) to condition 1 (common ROI-ROI analysis).

### Characterization of brain-states relevant to the cortical-depth

Dynamic analysis of layer-to-layer functional connectivity in healthy subjects distinguished two connectivity states, one characterized by intermediate-superficial layers being connected more strongly with other intermediate-superficial layers, and another where intermediate-deeper layers connected mainly with other intermediate-deeper layers, hereafter referred to as State One and State Two, respectively (**Fig. 3a**)). Nine out of twelve subjects spent most of the scan time in a state characterized by higher connectivity among the deeper cortical layers, but the difference in state duration was not significant (mean ± SD = 0.31±0.34 and 0.69±0.34 for State One and State Two respectively, N=12) (**Fig. 3b**). Although the main ROIs and layers involved in whole-brain connectivity were fairly similar across both states (**Fig. 3c**, top graphs), some differences were statistically significant (**Fig. 3c**, bottom graphs); however, these were individual connections, and no clear pattern was identified, with the exception of a potentially enhanced fronto-parietal and fronto-occipital connectivity in State One compared to State Two.

**Figure 3.**
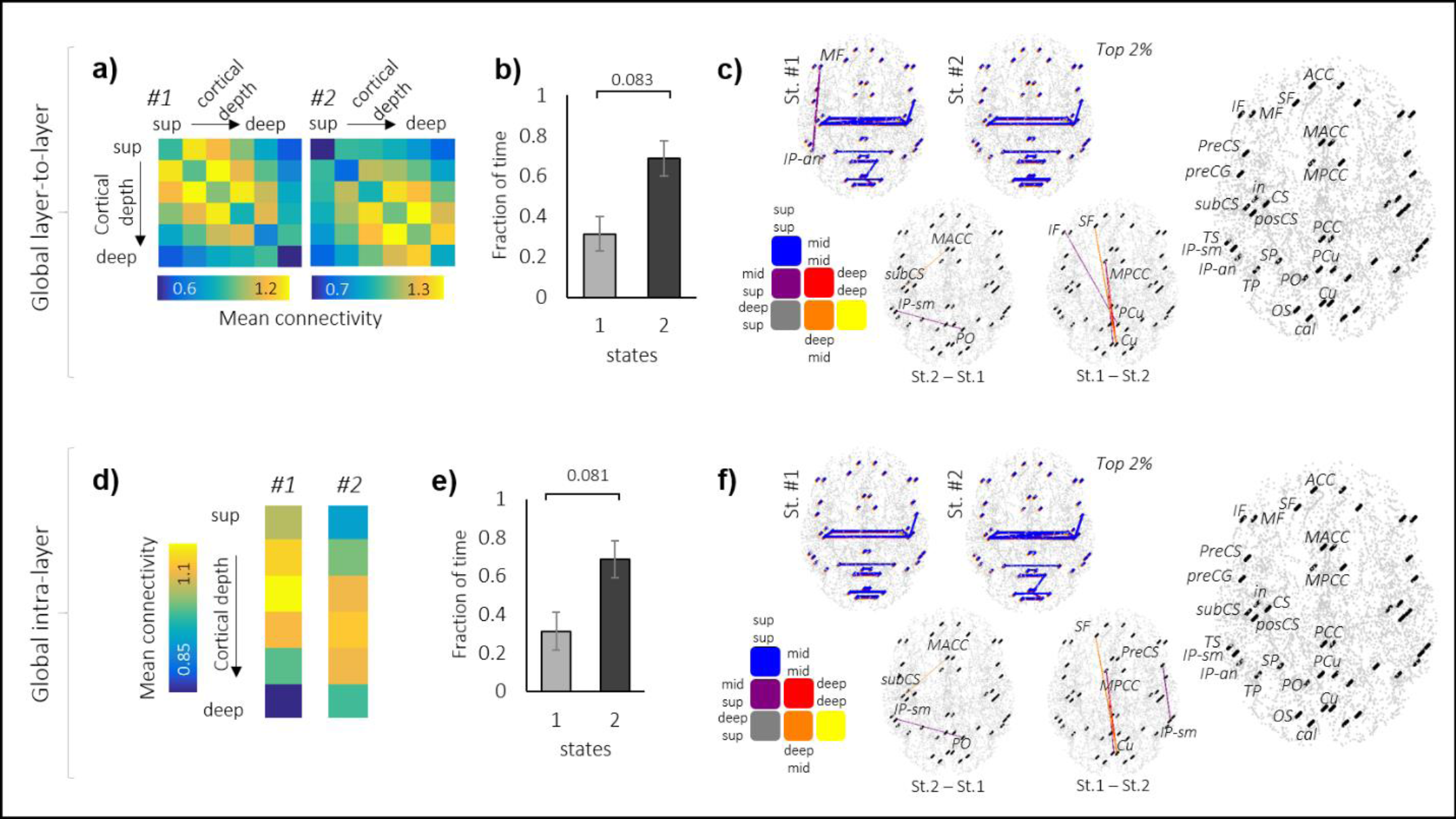
Cortical depth-based state identification. (**a**) Analysis of the time-varying layer-to-layer connectivity resulting in the identification of two different states, characterized by stronger connectivity among the intermediate-superficial layers (State One) or intermediate-deep layers (State Two). (**b**) Fraction of time spent in State One or Two (mean ± SE). The number over the horizontal bar indicates the p-value of a paired t-test (n=12). (**c**) Whole-cortex connectivity graphs associated with the identified brain states, colour-coded based on the layers involved. The top graphs show the top 2% connections associated with States One and Two. The graphs below show the differences between both states. All graphs are thresholded at a p-value <0.05 (corrected t-test). The graph on the right provides the ROI labels. (**d**) Analysis of the mean global intra-layer connectivity yielded two states resembling the intermediate-high and intermediate-low connectivity patterns observed in a). (**e-f**) same as b-c for states identified based on the global layer-to-layer connectivity. Note the remarkably similar results obtained with both inter-layer and global intra-layer analysis, which suggests that the target layers of laminar connectivity may not add critical information to differentiate cortical-depth connectivity states.

The global laminar analysis reflected similar results to the layer-to-layer analysis (**Fig. 3d-f**), i.e., the intermediate-superficial and the intermediate-deep portion of the cerebral cortex exhibited the strongest connectivity in State One and State Two, respectively, with similar prevalence and associated whole-brain connections as reported above.

### Triple network analysis along the cortical depth

In order to investigate network effects in the laminar dynamic functional connectivity analysis, we averaged the layer-to-layer connectivity values across the main ROIs that participate in the DMN, the SN and the CEN (**Table 1**). We concatenated the three connectivity matrices along the time dimension and performed k-means clustering to identify cortical states common to all three networks and the twelve healthy subjects (**Fig. 4a**). The calculated UI was 1.42±1.73 in the DMN, 3.00±2.63 in the CEN and 3.33±2.02 in the SN, and a paired t-test identified a significant difference between the DMN and the SN (p-value=0.008), suggesting that the cortical states were found to be significantly more stable (i.e., varied less) in the DMN compared to the SN (**Fig. 4b**, right panel). Cortical states were characterized in the three networks by intermediate-superficial or intermediate-deep laminar connectivity dominance in a similar way to the global analysis across the whole brain. State One, characterized by superficial connectivity, was significantly more dominant than State Two in the DMN (79.85±8.85 vs. 20.25±8.85, with p-value=0.004), while the prevalence of the two states was not significantly different in the SN or the CEN (**Fig. 4c**). The prevalence of both states in the DMN was significantly different from the prevalence of both states in the SN (p-value <0.05 in all cases); additionally, the prevalence of states in the DMN and their complementary in the CEN (e.g., state 1 of the DMN vs. state 2 of the CEN or vice versa) was significantly different (p-value=0.011).

**Figure 4.**
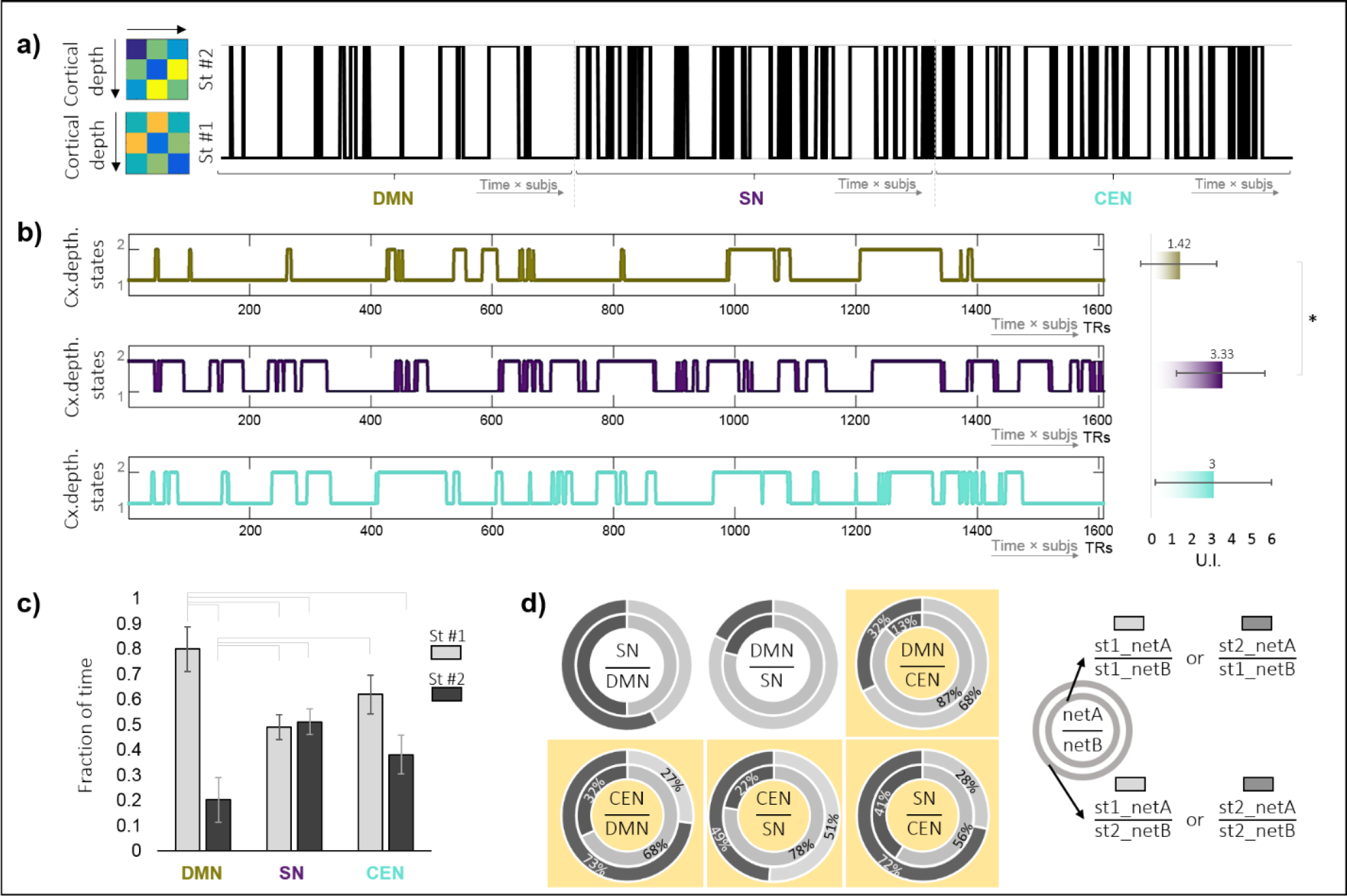
Cortical depth-based states in the triple network. (**a**) State fluctuation considering the depth-dependent connectivity matrices associated with DMN, SN and CEN simultaneously. State One and State Two (strongest connectivity in superficial and deep laminae, respectively) were identified in the three networks. The x-axis represents time points × subjects in the three cortical networks, concatenated. (**b**) State fluctuation as in **a** for each cortical network, and mean unrest index ± SD (right panel). (**c**) Fraction of time spent per state and network (mean ± SE, n=12 subjects). Horizontal bars indicate significant differences (p-value <0.05). (**d**) Relationship between States One and Two across the three networks (see legend on the right). The inner and outer circumferences represent State One and State Two of the network on the denominator, respectively. In each circumference, the portion shaded in light or dark grey indicates the proportion of time overlapping with State One or Two in the network of the numerator, respectively. E.g., State One of the CEN had a prevalence of 68% when the DMN was in State One but only 27% when the DMN was in State Two (**Table 2** provides numerical values). Yellow boxes highlight significant state prevalence changes that depend on the denominator network state. DMN: default mode network; SN: salience network; CEN: central executive control network.

**Table 1.** ROIs considered in the network analysis

| Network | ROIs (Destrieux atlas) |
| --- | --- |
| DMN | Precuneus |
|  | Middle anterior cingulate |
|  | Middle posterior cingulate |
|  | Posterior cingulate |
|  | Inferior parietal- angular |
| SN | Anterior cingulate |
|  | Insula |
| CEN | Superior frontal |
|  | Middle frontal |
|  | Inferior frontal |
|  | Inferior parietal – supramarginal |
|  | Inferior parietal- angular |

DMN: default mode network; SN: salience network; CEN: central executive control network.

**Table 2.**
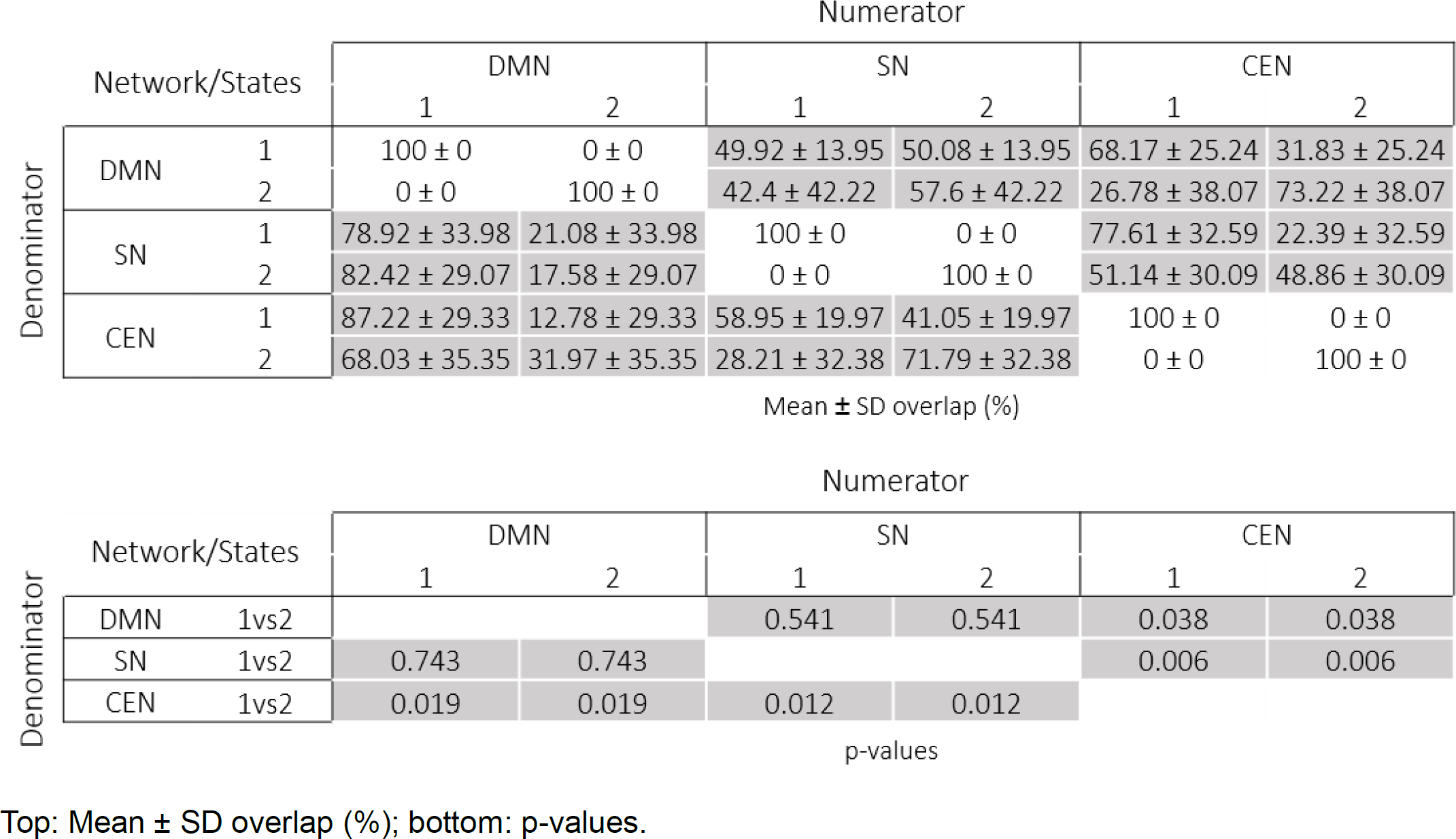
Relationship between State One and State Two across networks

We also evaluated potential network interplay by assessing the correspondence between states across networks. For each pair of networks (A and B), **Figure 4d** shows the percentage of time that State One (inner circle) or 2 (outer circle) of network B overlapped with State One (light grey) or 2 (dark grey) of network A. The corresponding numerical values are shown in **Table 2**. Independently of their cortical-depth state, the SN and the CEN were predominantly associated with the DMN in State One but not State Two (top middle and right panels in Fig. 4d, consistent with a more prevalent State One in the DMN). The SN transited between States One and Two approximately 50% of the time during either state of the DMN (top left panel in Fig. 4d). Significant differences were found concerning the state of the CEN depending on the DMN (bottom left panel in Fig. 4d), with a relatively strong agreement between the states in both networks (i.e., there was an average ∼68% and ∼73% overlap between State One and State Two of both networks, respectively). The CEN cortical state also varied significantly depending on the state of the SN (bottom middle panel in Fig. 4d), with ∼78% of the time in State One during State One of the SN but only ∼51% in this state when the SN was in State Two. Similarly, states 1 and 2 of the SN matched the corresponding states in the CEN ∼59% and ∼72% of the time, respectively (bottom right panel in Fig. 4d). The difference in the behaviour of the DMN depending on the CEN was also noted as significant (top right panel in Fig. 4d); although State One was always more prevalent in the DMN, its prevalence was ∼87%, where the CEN was also in State One, but only 68% when the CEN was in State Two. This cortical-depth state analysis involving the three networks provided an indirect way to study inter-network relationships and demonstrated common connectivity patterns concerning the depth of the connections involved in intra-network connectivity.

K-means identified eight different configurations according to the cortical-depth connectivity state of the three networks (**Fig. 5b**). Of these, the most prevalent was the one characterized by superficial connectivity in all three networks (cortical-depth State One in DMN, SN and CEN, triple network configuration 1, **Fig. 5b-c**), which appeared approximately one-third of the time across subjects). Approximately one-third of the time, the DMN and CEN were in cortical-depth State One while SN was in cortical-depth State Two. It was relatively uncommon for the SN and CEN networks to be in State One while DMN was in State Two (triple network configuration 7). The least common state was the one where the connectivity mode in the DMN and the SN involved deep layers of the cortex (cortical-depth State Two) while the CEN synchrony was driven by superficial connections (cortical-depth State One) (triple network configuration 8). This analysis brought new insights into the time-varying behaviour of resting-state network connectivity at a laminar level.

**Figure 5.**
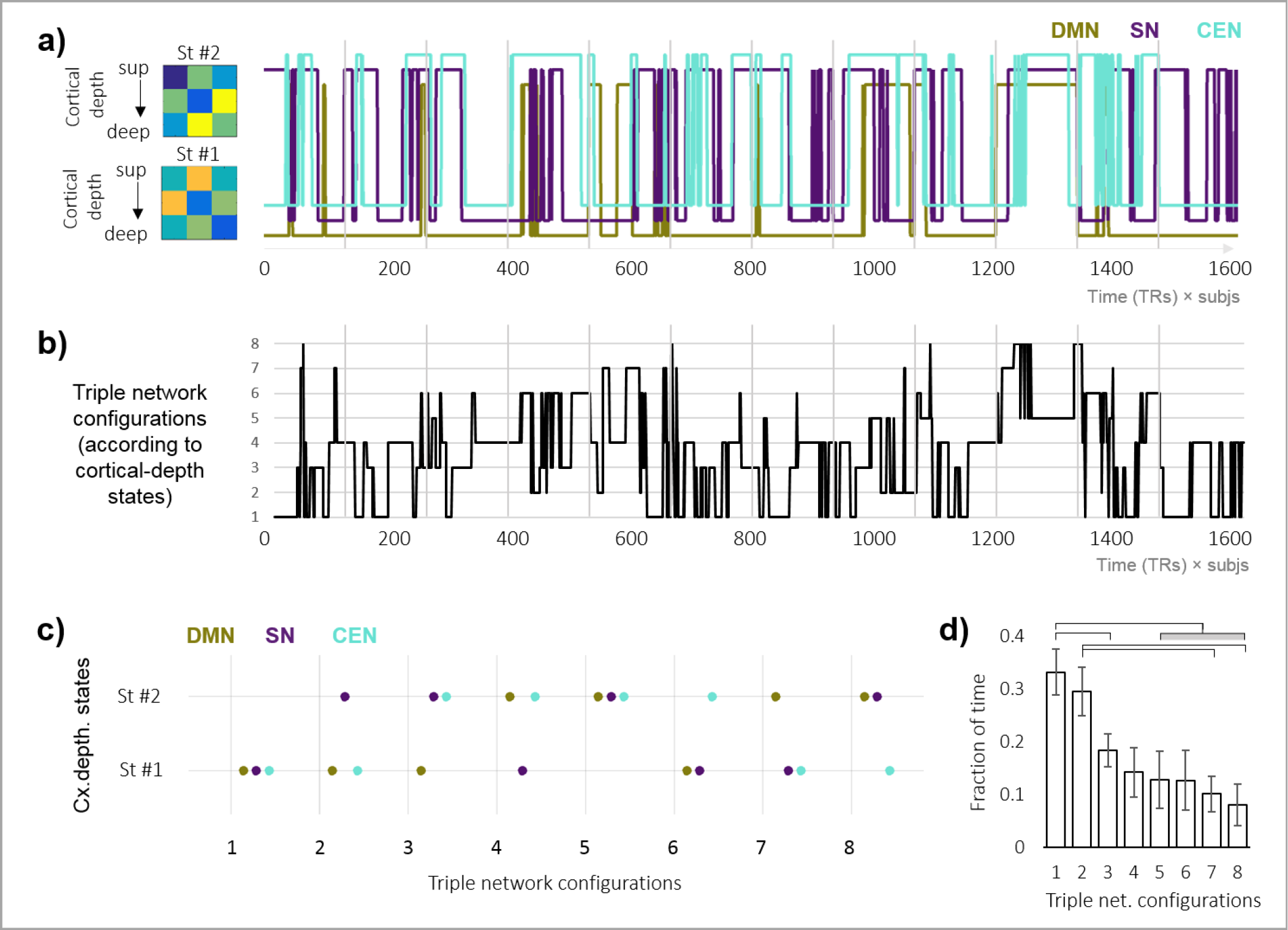
Triple network configuration states. (**a**) The three networks (each one in a different color) fluctuate in terms of cortical depth connectivity between a superficial and a deep connectivity state. This panel replicates Fig. 4b but with all the networks overlapping. (**b**) The graph shows the fluctuation of the triple network configurations, determined based on the co-occurrence of cortical-depth states for the DMN, the SN and the CEN (based on Fig. 5a). (**c**) According to state overlap among networks, eight different triple network configuration states were identified, shown in decreasing order of prevalence from left to right. E.g. configuration two was characterized by superficial dominance in the DMN and CEN (cortical-depth State One) but deep-layer dominance in SN (cortical-depth State Two). (**d**) Fraction of the time spent in each triple network configuration, the most common one corresponding to all three networks being in cortical-depth State One (triple network configuration 1). Horizontal lines connect significantly different groups (non-paired t-test due to unequal presence of states through subjects, p-value<0.05). DMN: default mode network; SN: salience network; CEN: central executive control network.

## DISCUSSION

In this work, we investigated the dynamic connectivity of the human cerebral cortex *in vivo* in healthy volunteers and demonstrated that a depth-dependent analysis can uncover dynamicity features of the human brain not trackable with typical ROI-based methods. Two highly dynamic cortical connectivity states were observed in a global laminar analysis, characterized by connectivity dominance of either more superficial or deeper cortical layers. Network analysis revealed that the superficial-connectivity state was predominant in the DMN and identified several cortical-depth dependent network configurations, suggesting that the cortical depth at which networks communicate can vary during rest.

### Dichotomy between superficial- and deep-layer connectivity states

Our k-means based analysis of global layer connectivity matrices clustered the data into superficial and deep connectivity states only, i.e., there was not a third state characterized by a central layer connectivity. This could be interpreted as a lack of feed-forward or bottom-up processing during resting-state (consistent with the absence of directed tasks); however, it should be borne in mind that, in our global analysis, the connectivity across all regions in the cerebral cortex was averaged, and hence, there is a possibility that, in some particular areas, feed-forward connectivity states may have ruled at certain times.

Although differences exist between particular subsystems, both the superficial and the deep layers of the cortex have been associated with feedback modulation across multiple brain areas, e.g., perceptual contour filling related to deep-layer activity in the visual cortex (Kok et al., 2016), auditory-based attention leading to superficial layer responses in the auditory cortex (De Martino et al., 2015), reading meaningful vs. non-meaningful words resulting in increased activity of deep layers in the left occipital temporal superior sulcus (Sharoh et al., 2019), responses to numerosity (perceiving the size of a group of items) in the parietal cortex strengthened in both, superficial and deep, but not in central layers (van Dijk et al., 2021). Diffusion-based studies have demonstrated a non-linear cortical-depth dependent myelination only in high-order brain areas, where deep and superficial layers showed a stronger myelination compared to the intermediate layers (Sui et al., 2021); this is in contrast to a linear trend (i.e., each layer exhibiting a larger level of myelination than the previous one) observed in primary unimodal areas (Sui et al., 2021).

In the visual system, superficial layers drive cortico-cortical connections, while deep layers are associated with subcortical modulation (Maier et al., 2010). Here, the deep layers are generally regarded as drivers of diffuse feedback processing and have been shown to receive input from the superficial layers, which exhibit connectivity patterns in the form of feed-forward but also feedback processing (Markov et al., 2014, Shushruth et al., 2009). While monocular stimulation leads to larger responses in deep and intermediate layers, it is the superficial laminae that seems more specific to spatial attention (de Hollander et al., 2021), demonstrating the complexity of the laminar visual system. Similar to the superficial-to-deep influence in the visual cortex, an optogenetics study showed that deep layers are modulated by superficial layers in primary somatosensory areas (inhibited by means of interneurons) (Pluta et al., 2019). In the auditory system, the complexity of responses observed at superficial and deep layers compared to intermediate territories has been hypothesized to relate to columnar processing, horizontal connections and cortico-cortictal feedback (Moerel et al., 2019). High order areas such as the prefrontal cortex are strongly connected with supragranular but not infragranular layers according to fiber tracer studies (Kritzer and Goldman-Rakic, 1995), and multiple frontal, parietal and temporal areas functionally connect through superficial layers, which are believed to contribute to the slow rhythms of EEG during wakefulness and sleep (Halgren et al., 2018). In agreement, supragranular layers but not infragranular layers seem to be responsible for the transfer of information between DMN areas (Whitesell et al., 2021). Although the meaning of global superficial-layer and global deep-layer connectivity remains unknown, our results suggest that these states, both probably related to high-order processing, constitute different working modes of the brain, potentially affected by different sub-systems during quiet wakefulness.

### Selected fMRI sequence

For this study, we made use of an fMRI sequence (EPIK) that is based on GE-EPI but combines a series of acceleration techniques and alternate peripheral k-space sampling to offer higher spatial resolution and lower image distortions compared to the standard method (Yun et al., 2013, Yun and Shah, 2017, Caldeira et al., 2019, Shah et al., 2019, Yun et al., 2019, Shah, 2003, Shah NJ, 2004, Zaitsev et al., 2001, Zaitsev et al., 2005, Yun and Shah, 2020, Yun et al., 2022). Given the gradient-echo BOLD nature of the sequence and its potential bias towards large veins, we have devoted recent publications to validate its use in laminar applications (Pais-Roldán et al., 2023) (other examples of GE-BOLD applied to laminar fMRI in particular sub-systems can be found in the literature, e.g. (Siero et al., 2015, Havlicek and Uludag, 2020, Kay et al., 2019)). Some activity measures derived from this kind of high-resolution fMRI data are possibly affected by a vein-related superficial bias, e.g. the amplitude of low-frequency fluctuations may be systematically stronger in superficial cortical layers; however, the connectivity patterns along the cortical depth assessed with EPIK vary across different brain networks and during task performance (Pais-Roldán et al., 2023), which indicates that the high-resolution method employed is able to track laminar differences relevant to brain function. Lastly, the implementation of EPIK used here allows whole-brain coverage together with a spatial resolution sufficient to explore cortical depth (0.63 × 0.63 × 0.63mm) and a moderate TR of 3.5s. These three conditions together have allowed us to explore whole-brain resting-state networks and their cortical depth dependence.

### Dynamicity of connections along the cortical depth

Before considering why the laminar connectivity is more susceptible to change than the ROI-ROI communication, it is important to first rule out potential contamination of the laminar data. Here, we have investigated the motion content in each sliding window as the main parameter that could affect our laminar analysis and found no correlation between the identified state switches and the amount of motion. Assuming the laminar fluctuations of connectivity are true, it appears that independently of the ROI-ROI correlations, the interconnected mesh below the cortical surface is not hard-wired; instead, different moments coincide with a higher or lower degree of connectivity of the different depth-territories. In a previous report (Pais-Roldán et al., 2023), we described the laminar features of diverse brain networks, and we observed several differences, at a whole-brain level, in the laminar function during rest and during task. The present study complements our previous results by showing that laminar changes do not only pertain to sufficiently different states, e.g. rest vs. task, but they can also be observed in successive periods of the same resting-state scan. Although the complex interplay between the layers of remote cortical areas is not yet well understood, given the clarity of our results, we hypothesize that this cortical dimension (depth) plays a critical role in the maintenance of the brain state and, as such, it could constitute a promising target to investigate brain state alterations, a topic that we are investigating in parallel studies.

One question that remains is why the analysis of a high dimensionality matrix containing both ROI and cortical-depth information did not result in at least the same number of state switches as those identified from a layer-only-based connectivity matrix. It is possible that the large amount of information fed to the k-means algorithm obscured the global effects taking place along the cortical thickness, as more data usually means higher variability and more difficulty to find clusters of low intra-cluster variance.

### Laminar dynamic connectivity in the triple network

Besides the global whole-brain analysis, our network analysis reported laminar differences, both in terms of state variability and preferred layer of communication. Considering the whole brain, a deep-layer communication was predominant within the scan time for most subjects; however, this was not the case for the analysis that focused on the DMN. The higher prevalence of the superficial-connectivity state in the DMN is in agreement with our previous static connectivity results, where the DMN showed maximum correlation among intermediate-superficial layers (Pais-Roldán et al., 2023). Animal studies also suggest that layers II-III play a critical role in the maintenance of DMN communication (Whitesell et al., 2021). The fact that connections in the DMN nodes are predominantly, but not exclusively, superficial poses the question of what makes the preferred depth of communication change. To answer this question, high-resolution evaluations with added disruption/stimulation paradigms or with an external evaluation of the brain state to cross-reference with the laminar working modes may be necessary.

Although the network that has been investigated the most during resting-state is the DMN, which is involved in mental processes that take place mostly in the absence of a task (Raichle et al., 2001), its interplay with the CEN and the SN, i.e., the “triple network” (Menon, 2011) has recently gained attention due to its potential role in psychopathology and mind-shaping (Menon, 2019, Altinok et al., 2021). According to the triple network hypothesis, the SN acts as a switch between the DMN and CEN (Kronke et al., 2020, Nekovarova et al., 2014, Whitesell et al., 2021). Our results show that most of the time, the layers involved in the connectivity of the three networks are superficial. However, it was also common for the SN to use deeper territories to communicate and, somewhat less often, this was true for the CEN too (in this situation, the DMN was the only network whose connectivity was persistently driven by superficial layers). The DMN was also the network where states fluctuated the least, while the SN was significantly more dynamic. Interestingly, most paired-network combinations showed a significant co-dependence on their cortical-depth state. These findings have three main implications. First, all cortical territories are accessible by all three network systems, i.e., although a network may have a laminar preference, the main laminar hub of communication can change even during the course of one resting-state scan. Second, the DMN seems significantly more stable in terms of laminar preference than other networks (it showed a lower unrest index). The laminar stability of the DMN observed here is in agreement with axonal tracing studies in mice showing that the superficial layers of cortical areas of the DMN project almost exclusively to other DMN territories. This is in contrast with fibres starting at deeper layers of the DMN, which have diverse projections (Whitesell et al., 2021). Third, the fact that the SN showed the highest cortical-depth state variability would support a hypothetical role of the cortical depth in network switching, e.g., the SN could employ superficial or deeper layers to communicate with the DMN or the CEN, respectively. Although future studies should further investigate the laminar implications of triple network connectivity, the similarities and differences found here pertaining to the three networks in terms of the cortical-depths involved and the unrest-index of this laminar connectivity are aspects of the triple network configuration that may be of value for understanding neuro-psychiatric disorders.

### Static vs. dynamic functional connectivity

Some literature suggests that the analysis of static connectivity could identify specific features of a subject better than time-varying connectivity measures (Menon and Krishnamurthy, 2019). While we did not observe radical state changes involving ROI-ROI connectivity in our subjects, the dynamic connectivity analysis performed on laminar connectivity matrices revealed highly variable states. Although the meaning of this variable strength of cortical connectivity requires better understanding, we believe that static and dynamic connectivity can be complementary parts of a functional analysis, the first one providing more robust features of the system and the second one being more attractive to investigate brain state fluctuation. In particular, we have shown that connectivity along the cortical depth seems to be much more variable than across the cortical surface, suggesting that a dynamic analysis of the laminar connectivity could add critical information when monitoring brain states.

### Limitations

The high-dimensional k-means clustering performed here for state identification might be hampered by the relatively low sample size and wide age-range employed in this study. Additionally, although we demonstrate an impact of using cortical depth information on state identification, we have not verified the relevance of the identified cortical states. This could be assessed, for instance, by applying the cortical depth-dependent dynamic functional connectivity analysis to high-resolution data from a patient population compared to a healthy control group.

Here we have shown how dynamic functional connectivity not only applies to the varying relationship between distant ROIs but also to the functional organization of the cortical layers in the healthy human brain, which suggests that this dimension of the cortex contributes to the varying brain state during wakefulness. Future studies will investigate the relevance of these laminar connectivity switches in the context of neuropsychiatric disorders.

## MATERIALS AND METHODS

### Experimental Design

The main objective of the study was to assess the potential benefit, in terms of brain state identification, of using a dynamic connectivity analysis focused on the cortical depth compared to the typical analysis involving cortical segmentation into ROIs. For this purpose, resting-state fMRI scans were acquired with high spatial resolution and were analysed using different sets of information (a total of 26 conditions involving different combinations of ROIs and cortical-depths were evaluated, see section “Dynamic Functional Connectivity Analysis”). State identification using the cortical-depth based analysis was followed by state characterization (see “Analysis of Brain States”) and network-based analysis (see “State Identification in the triple Network” and “Network Interaction”). The specific methods employed are described below.

### Subjects and Data Acquisition

The data from twelve healthy subjects (nine males and three females, 29.5±6.2 years) were used in the present study. The experimental methods were approved by the local institutional review board (EK 346/17, RWTH Aachen University, Germany), MR-safety screening was performed prior to MRI acquisition, and informed written consent was obtained from all subjects. All data were collected using a Siemens Magnetom Terra 7T scanner with a 1-channel Tx / 32-channel Rx Nova Medical head coil. One resting-state fMRI scan (172 volumes; ∼10min) was obtained from each subject using a gradient-echo EPIK sequence (Echo Planar Imaging with Keyhole) with the following parameters: TR/TE = 3500/22 ms, FA = 85°, partial Fourier = 5/8, 3-fold in-plane/3-fold inter-plane (multi-band) acceleration, matrix = 336 × 336 × 123 slices (voxel size: 0.63 × 0.63 × 0.63 mm3). Subjects were instructed to keep their eyes closed, not to sleep and not to think about anything in particular. B0 shimming was performed with a standard routine provided by the manufacturer. A structural scan was also obtained, for tissue segmentation purposes, using an MP2RAGE sequence with TR/TE = 4300/2 ms, matrix = 256 × 376 × 400 (voxel size: 0.60 × 0.60 × 0.60 mm3). Pulse and breathing were monitored during the acquisition using a finger oximeter and a respiratory belt for post-hoc fMRI de-noising.

### Data Pre-processing

The functional images, both magnitude and phase, were slice-timing corrected (SPM12, Statistical Parametric Mapping Software, UCL, London, UK), the first four TRs were removed, and the remaining images were realigned (AFNI, Analysis of Functional NeuroImages, NIH, Bethesda, MD), band-pass filtered (0.005-0.12Hz, AFNI) and de-noised (AFNI) by regressing out the mean signal of the CSF and white matter tissue, the motion parameters (three demeaned and three derivative regressors), and the physiological parameters (generated by estimating Fourier sine and cosine series fit of the cardiac and respiratory signals, plus multiplicative terms: https://github.com/tesswallace/retroicor/blob/master/mod_retroicor.m). A de-veining step, consisting of subtracting a fitted version of the functional phase image from the functional magnitude image, was added to the pre-processing pipeline, as previously described (Menon, 2002, Curtis et al., 2014, Pais-Roldán et al., 2023). The anatomical scan was manually co-registered to the functional volume for each subject, and a mask was manually drawn over imperfect co-registration areas to exclude those voxels from the analysis (manual co-registration and masking were done based on visual inspection on Freeview (Freesurfer, Martinos Center for Biomedical Imaging, Charlestown, MA)). Segmentation of the co-registered anatomical data was performed with Freesurfer using recon-all. The segmentation of the cerebral cortex into different areas was based on the Destrieux cortical atlas (Destrieux et al., 2010, Duvernoy, 1999). The cortex was additionally segmented into six tangential territories (a proxy for cortical depth). Briefly, the cortical ribbon, i.e., the space between the outer and the inner surface of the cortex, was split into six surfaces, each separated by the same distance, using mris_expand. Voxels in the functional image were projected to the vertices of the six cortical surfaces through trilinear interpolation using the function mri_vol2surf on Freesurfer. The resulting functional surface was then subjected to 1 mm within-surface smoothing.

### Dynamic Functional Connectivity Analysis

The mean time course of 46 cortical ROIs (23 ROIs selected from the cortical atlas in each cerebral hemisphere) at six cortical depths were extracted using Matlab R2015a (Mathworks, Natick, MA, USA), and the correlation between each pair of time courses was computed using sliding windows of length 120s and step size 3.5s (1TR). This resulted in 134 connectivity matrices per resting-state scan. In order to make the analysis sensitive to ROI/layer connectivity changes and not the overall level of brain connectivity, each sliding-window connectivity matrix was normalized by dividing each value by the average value of the sliding window. Twenty-six types of dynamic connectivity analyses were performed to evaluate how different segmentation of the cortex (connections evaluated at different levels) could affect the identification of brain states (see **Fig. 1** and **Fig. 2**). The twenty-six conditions were: 1. Ignoring the depth-dimension of the cortex (normal ROI-connectivity analysis with matrix size=46×46, **Fig. 1b**); 2-7. Evaluating the connections between all ROIs within each of the six cortical depths individually (matrix size=46×46, **Fig. 1c**); 8-22. Evaluating all connections between the ROIs at cortical depth M and all the ROIs at cortical depth N (matrix size= 46×46, **Fig. 1d**); 23. Considering all layers of all ROIs (matrix size=276×276, **Fig. 1a**); 24. Considering all layers of all ROIs except for intra-layer connections (matrix size=276×276, with 46 NANs per row, **Fig. 1e**); 25. Narrowing the analysis to the layer-to-layer global connectivity (matrix size=6×6, **Fig. 1f**); 26. Further condensing information into a mean intra-layer connectivity vector (matrix size=6×1, **Fig. 1g**). Auto-correlations (matrix diagonal in symmetric matrices) were excluded (substituted by NANs). We refer to each one of these analysis as a different “condition”, “analysis scheme” or “pipeline” along the text.

### State Decomposition

To identify states common to all subjects, the 134 connectivity matrices of all 12 subjects were first triangulated (to avoid symmetric repetitions), reshaped into vectors of length “*VLength*” and concatenated along the time domain. The resulting input-matrices, of size *[VLength × (134×12)]* were subsequently fed into a k-means algorithm to automatically cluster the matrices by minimizing the intra-group variance. We used a Matlab implementation of k-means, the Calinski-Harabasz index (variance ratio criterion), and a range of 1-20 clusters. The optimal number of clusters was between two and four for all conditions. The result of the k-means analysis was a clustering-vector of size *[1 × 134×12]* (1608 time points) where the value of each cell indicated the state associated with that particular time point (e.g. “1” or “2”, in the case where an optimal number of states is equal to 2).

### Motion Content

To evaluate a potential relationship between subject motion and the dynamic connectivity results, the motion content was calculated in sliding windows by matching the dynamic connectivity analysis using the following formula:

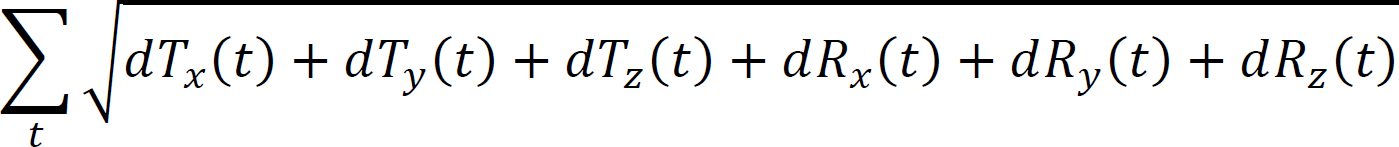

, where T_x_, T_y_, T_z_ and R_x_, R_y_, R_z_ are translational and rotational motion regressors retrieved during realignment, and t is the duration (in TRs) of the sliding window.

### Analysis of Brain States

An unrest index (UI) was computed as the mean number of state switches per scan and subject for each one of the 26 analysis conditions. This allowed us to quantify the volatility of the states and of the different connections during a resting state and to determine, for instance, whether the ROI-ROI connections were more or less dynamic than the global connectivity between cortical laminae.

The specific pattern of connectivity associated with each particular state was computed by averaging the connectivity matrices of all the time points assigned to that particular state, e.g., in a 2-states scenario, the first dimension of the input-matrices (space) was averaged across the second dimension (time) at indexes that corresponded to State One or State Two in the clustering-vector.

To quantify the dwell time of the identified states, the clustering-vector was reshaped into a *[134 × 12]* matrix (i.e., converted from a concatenated-subject to subject-specific time vectors), and the mean time spent per state was computed by averaging across all subjects the number of cells matching the specific state.

To compute the whole-brain connectivity patterns associated with the states identified based on layers only, the matrices in pipeline 23 (all ROIs, all layers) were averaged following the clustering-vector of analysis pipelines 25 and 26. This was first done at a subject level (i.e., for each of the 12 subjects), and the subject-specific matrices were then subjected to statistical group analysis. A one-sample t-test was computed for each cell of the matrices, and the mean correlations were thresholded by p-value (0.05, corrected using the fdr_bh function in matlab) and mean correlation value (calculated as the value corresponding to a 98% percentile - top 2% connections). Differences between states were assessed with a paired t-test. Whole-brain graphs were generated using a customized Matlab code, making use of the information in the Freesurfer file aparc2009_annot (cortical atlas in 3D-surface space).

### State Identification in the Triple Network

In order to investigate cortical-depth states at a network level, we performed k-means clustering of layer-to-layer connectivity matrices computed from all ROIs involved in three brain networks: default mode (DMN), salience (SN) and central executive control (CEN), which are often referred to as the triple network (Menon, 2011) (the ROIs used to compute the mean connectivity matrix are provided on **Table 1**). Given the smooth pattern of the layer-based results (i.e., a superficial or a deep layer gradient), the layer-to-layer input connectivity matrix used for this analysis was a simplification of the 6 by 6 matrix, where layers were averaged in pairs to yield a 3 by 3 matrix (only three cortical territories: superficial, intermediate and deep, for straightforwardness). To identify cortical-states common to all three networks, the three space-time input-matrices were further concatenated, and k-means was performed on a new input-matrix of size *[6 × 134*12*3]*.

The algorithm returned a vector of size *[1 × 134*12*3]*, which could be reshaped into a matrix of size *[3 × 134*12]* to track network-specific results, with two main states that oscillated in time for all three networks. The data were then subjected to UI determination, state-duration analysis and state overlap analysis (network state synchrony –see next-).

### Network Interaction (State Overlap Analysis)

A proxy of network communication was estimated based on network state synchrony, i.e., overlap of the cortical-depth state between networks, calculated as the percentage of the time that a network state was shared with other networks. To detect the different configurations of the triple network according to the cortical-depth state, k-means was performed again on this data (the clustering-vector of the three networks in time, of size = *[3 × 134*12]*), which returned eight different configurations or triple network states. Finally, the time spent in each configuration was computed in the same way as previously described.

### Statistical Analysis

Paired t-tests were performed to assess the significance of the differences in UI detected between conditions as well as the differences in prevalence between the two identified states at whole-brain and at network level. A one-sample t-test was performed to detect significant connections within each cortical state. A two-sample unpaired t-test was performed to assess the statistical significance of the difference in prevalence between each pair of network configurations, as not all configurations were observed in all subjects. All the analyses performed here refer to the same group of subjects (N=12). Results were considered significant if the obtained p-value was below 0.05.

## ACKNOWLEDGEMENTS

We thank Ms Elke Bechholz and Ms Anita Köth for technical support during MRI acquisition; Ms Rick Claire for manuscript corrections, and all the volunteers for their excellent cooperation.

## Funding

The authors acknowledge that they have received internal funding from the Forschungszentrum Juelich (Helmholz association) for this research.

## Author contributions

N.J.S. designed the research; S.Y. developed the MR imaging sequence and the corresponding image reconstruction software; S.Y. and P.P. performed in vivo experiments. P.P. and S.R. designed and performed data analysis; M.Z. and J.F. developed and maintained computer resources to pre-process the data, P.P. wrote the manuscript and prepared the figures; N.J.S. invented the original EPIK sequence; S.Y., S.R., T.V., I.N. and N.J.S. reviewed the manuscript.

## Competing interests

The authors declare that they have no competing interests.

## Data and materials availability

All data needed to evaluate the conclusions on the paper are present in the paper. The MRI datasets can be shared upon formal request to <u>.</u>

## Notes

### Competing Interest Statement

The authors have declared no competing interest.

