## Supplementary Text for "Contribution of cortical layers to human dynamic functional connectivity"

**This PDF file includes:**

Supplementary Text

### Supplementary Text

**Conditions tested in the initial dynamic functional connectivity analysis: implications of matrix content on the resolution of brain states (relevant to Fig. 2)**

After computing a simple ROI-ROI analysis (condition 1), we considered the data on an ROI-ROI basis but focused on a particular cortical depth to assess whether one of the six cortical representations drives the whole resting-state connectivity (conditions 2-7). Of note, all matrices were normalized at each time point to make the clustering procedure sensitive to changes among ROIs/layers and not absolute connectivity values. The relevance of a layer in state determination was quantified as the number of state switches (brain unrest index) observed when the analysis was based on that layer's connections. All cortical depths assessed individually resulted in similar brain-state dynamics, suggesting that no layer alone was responsible for brain dynamicity. We also performed an analysis considering correlations between the ROIs of one layer and the ROIs of a different layer (conditions 8-22). Evaluation of the unrest index associated with the analysis of all pair combinations suggested that the connections between layers 2 and 4, and between layers 3 and 4 constituted a relatively important feature to sort the whole-brain connectivity matrices into different states. Considering all ROIs and all cortical depths, with or without exclusion of within-layer ROI-ROI connections (conditions 23 and 24, respectively), did not yield higher dynamicity compared to the common whole-cortical-ribbon ROI-ROI dynamic connectivity analysis. We then contemplated the possibility that the connectivity state of the cortical layers as a whole could vary over time, possibly reflecting more dynamic brain states than the ROI-ROI connections. To evaluate this, we averaged the connectivity values across ROIs on a layer-to-layer (condition 25) or a global intra-layer (condition 26) scheme, in a way that we could assess what cortical depth was more connected (and with which other layer, in the case of the layer-to-layers analysis) at each moment, disregarding the coordinates on the cortical surface. Compared to any other scenario, the analyses of connectivity purely based on layer information (conditions 25 and 26) resulted in the highest level of dynamicity, suggesting that the cortical depth could play an important role in the determination of the brain state.
